# Construction of a breast cancer predictive nomogram based on diverse cell death methods and reveal tumor microenvironment characterization

**DOI:** 10.1101/2024.05.09.593462

**Authors:** Rihan Wu, Zirui Wang, Chunhui Dong, Yihui Liu, Ling Chen

**Author notes:** Corresponding author: Ling Chen, 277 West Yanta Road, The First Affiliated Hospital of Xi’an Jiaotong University, Yanta District, Xi’an, Shaanxi Province, China 710061.

## Abstract

**Objective:** To develop a robust predictive model and nomogram for breast cancer (BC) linked to genes associated with diverse cell death methods.

**Methods:** The prognostic model was constructed using the LASSO Cox method. Model performance was assessed using K-M analysis, ROC curves, and independent prognostic analysis. Subsequently, we constructed a nomogram and analyzed differences in tumor microenvironment and drug sensitivity between different subgroups. The enrichment of differentially expressed genes between different subgroups was assessed. Additionally, we overexpressed CD24 in BC cell lines to assess its impact on cellular proliferation using CCK8 assays, migration through scratch and transwell assays, and apoptosis via flow cytometry.

**Results:** A prognostic model comprising twelve genes (CREB3L1, SFRP1, SHARPIN, AIFM1, IL-18, CD24, EDA2R, CRIP1, XBP1, BCL2A1, NKX3-1, and NME5) was constructed. BC patients were categorized into different subgroups, with the low-risk subgroup demonstrating superior survival. Additionally, we constructed a nomogram. The nomogram was validated as a reliable independent predictor of outcome. The enrichment analyses imply a connection between patient risk and immune response. The low-risk subgroup had a higher TME score. Patients in the high-risk group had improved responses to lapatinib, BI-2536, OSI-027, and SB505124, while those in the low-risk subgroup showed improved sensitivity to axitinib, epirubicin, fulvestrant, and olaparib. Furthermore, CD24 overexpression was found to promote proliferation and migration, while inhibiting apoptosis.

**Conclusion:** These findings contribute to the individualization of treatment and aid in uncovering the tumor microenvironment characterization for BC patients.

## Introduction

Based on the most recent data available from the American Cancer Society, in 2023, there were approximately 300,590 new cases of breast cancer (BC) and 43,700 BC-related fatalities among American women. BC is the main cause of death from cancer in women, second only to lung cancer, accounting for 31% of all newly diagnosed cases. The incidence of female BC has been slowly increasing by about 0.5% annually since the mid-2000s, posing a significant threat to women’s lives and health[1]. BC is characterized by high heterogeneity, presenting different pathological features, molecular subtypes, diverse treatment methods, and varying clinical prognoses[2]. Existing classification systems fall short of meeting the demand for precise cancer treatment. As a result, several molecular tests, including the 21-gene and 70-gene prediction models, have been developed for better predicting the long-term outcomes and therapeutic responses of BC patients[3–6]. Therefore, gaining a profound understanding of the pathogenesis, biological characteristics, and exploration of genetic profiles is essential for establishing effective personalized treatment strategies.

Apoptosis, the earliest and most well-known programmed cell death (PCD) method, involves a series of morphological changes, including chromatin condensation, cell shrinkage, cell membrane blebbing, and mitochondrial swelling[7, 8]. Pyroptosis, on the other hand, is typified by the forming of membrane pores, leading to cell enlargement until membrane rupture, which spills cellular contents and triggers a potent inflammatory response due to osmotic pressure changes[9]. Research indicates that pyroptosis might have a dual function in carcinogenesis. Some scholars argue that high GSDMB expression in BC cells promotes BC growth, invasion, and metastasis, contributing to high metastasis rates and lower survival rates in patients[10–12]. However, pyroptosis can also inhibit tumor initiation and progression. Studies by Xia Wu et al. have confirmed that BC patients with higher expression levels of GSDMD, IL-1β, and caspase-1 exhibit lower pathological grades and reduced metastatic potential[13]. Necroptosis, mediated by RIP1 and RIP3, is characterized by the massive release of cytoplasmic contents and organelle swelling[14–16]. Many studies have demonstrated the anti-tumor effect of necroptosis, although most tumor cells tend to evade this pathway[17, 18]. Ferroptosis, initiated by iron-dependent phospholipid peroxidation, is controlled by a number of cellular metabolic processes, including iron metabolism, redox homeostasis, mitochondrial function, and several disease-related signaling pathways[19]. Cuproptosis, a recently discovered mechanism, relies on copper-dependent cell death and is implicated in the pathophysiology of various illnesses[20, 21]. Studies have shown that patients with different cancers often exhibit elevated copper levels in tumor tissue and altered systemic copper distribution[22]. Disulfidptosis, a novel form of cell death, was reported and named by Professor Gan Boyi’s team in February 2023. It occurs under glucose-starvation conditions when there is an inadequate supply of nicotinamide adenine dinucleotide phosphate. Cancer cells that overexpress solute carrier family 7 member 11 accumulate disulfides, leading to cell death[23]. It is evident that these programmed cell death patterns are closely related to tumor initiation and development. However, the relationship between these cell death patterns and BC is not yet fully understood.

In our study, we identified survival-related differentially expressed genes (DEGs) of BC patients and developed a model and nomogram. Furthermore, we explored differences in immune microenvironments and medication responsiveness between patients in different risk subgroups. This finding will contribute to the precise individualized treatment of BC patients and provide insights into the impact of the tumor microenvironment on tumor initiation and progression.

## Materials and methods

### Data sources

The TCGA and the GEO database were used to extract the clinical information and gene expression data of BC patients. Inclusion criteria for dataset selection were as follows: (1) pathologically confirmed BC samples and para-carcinoma samples; (2) patients with complete survival data. Exclusion criteria comprised: (1) patients with 0 overall survival; (2) repeated samples of the same patient. Our dataset comprised RNA sequencing data and clinical information from 1058 BC samples and 99 para-carcinoma samples sourced from TCGA, along with an additional 327 BC samples from the GSE20685 dataset for our research. Diverse PCD-related genes were extracted from prior literature[24, 25]. Finally, a total of 765 genes were obtained.

### Identification of DEGs

We performed a differential analysis of raw transcriptome count data using the “edgeR” package in R for 1058 BC samples and 99 para-carcinoma tissue samples from the TCGA dataset. False Discovery Rate (FDR) < 0.05 and |logFC| > 0.585 were the criteria used to screen DEGs. The Search Tool for the Retrieval of Interacting Genes (STRING v12.0) was used to build a protein-protein interaction (PPI) network for the DEGs.

### Univariate Cox analysis

We combined the FPKM data from 1058 BC samples in the TCGA dataset with 327 BC samples from the GSE20685 dataset and removed the batch effect. The FPKM values of RNA sequencing data were converted into log data for subsequent analysis. Next, we employed univariate Cox analysis to identify prognosis-related DEGs. A significance threshold of less than 0.03 was applied for the P-value.

### Construction and validation of the predictive model

We initially randomized the dataset, consisting of 1385 BC patients, into training and testing cohorts in a 1:1 ratio using the “caret” package. To mitigate potential overfitting issues and create a score signature, we then applied the least absolute shrinkage and selection operator (LASSO) Cox regression analysis. Subsequently, the risk score (RS) for each sample was then calculated using the formula RS= Σ (Coefi * Expi), in which “Coefi” stands for coefficient and “Expi” for gene expression levels. Within the training cohort, we divided the samples into different risk subgroups according to the median RS. Kaplan-Meier (K-M) analysis was used to assess overall survival (OS). The ROC analysis and area under the curve (AUC) calculation were performed using the “timeROC” package. The same methodology was applied to the testing cohort. The multifactor Cox analysis was performed along with other clinicopathological factors to confirm the independent predictive effectiveness of the RS.

### Nomogram construction and calibration

We integrated the RS and relevant clinicopathologic factors to create a prognostic nomogram. To assess the nomogram’s performance, we generated a calibration plot that compared the predicted OS with the observed OS. Cumulative hazard curves were plotted using the “survminer” R package. Decision Curve Analysis (DCA) is a valuable tool for evaluating the clinical usefulness and benefits of a predictive model. We utilized the “devtools,” “survival,” “survminer,” and “ggDCA” packages to perform DCA.

### Enrichment analysis of the DEGs between different risk subgroups

We used the selection criterion of FDR < 0.05 and |logFC| ≥ 0.585 to identify DEGs between various risk subgroups. Subsequently, we conducted Gene Ontology (GO) and Kyoto Encyclopedia of Genes and Genomes (KEGG) pathway enrichment analyses. These analyses allowed us to discover the functional categories and biological pathways connected to the identified DEGs. Additionally, we used the “clusterProfiler” package to carry out gene set enrichment analysis (GSEA).

### Analysis of immune cell infiltration and TME

Using the “preprocessCore”, “e1071”, and “limma” packages, we computed the relative content of 22 immune cells to assess the extent of immune cell infiltration. To delve deeper into the variations in the fraction of immune cell types between different RS subgroups, we applied the CIBERSORT algorithm. Then, we analyzed the correlation of different immune cells with RS and selected genes used to build the model.

Additionally, the proportions of stromal and immune cells in different groups were computed utilizing the “utils”, “limma”, and “estimate” packages, deriving TME scores (including ESTIMATE scores, stromal scores, and immune scores). Finally, we visualized the findings using a violin plot generated with the “ggpubr” package.

### Drug sensitivity analysis

We examined variations in the therapeutic responses to drugs among BC patients in different RS subgroups. The semi-inhibitory concentration (IC50) of anticancer medications, utilized in the treatment of tumor patients, was computed using the “oncoPredict” package. Lower IC50 values suggest greater patient sensitivity to the drug.

### Plasmid Transfection in MCF7 Cells

MCF7 cells were seeded and allowed to grow for 24 hours until 70-90% confluency was reached. For transfection, 4 µg of plasmid DNA was diluted in 250 µL of DMEM without serum to prepare the DNA dilution. Separately, 10 µL of Lipofectamine 2000 was gently mixed with 250 µL of DMEM and incubated at room temperature for 5 minutes to form the Lipofectamine dilution. The two solutions were then combined and incubated at room temperature for 20 minutes to allow DNA-Lipofectamine complex formation. The medium from the cells in a 6-well plate was aspirated, and the DNA-Lipofectamine mixture was added directly to the cells. The plate was gently rocked back and forth to distribute the complexes evenly and then supplemented with 1.5 mL of fresh DMEM. After 5 hours of incubation at 37°C, the medium was replaced with complete growth medium, and the cells were further cultured for 72 hours.

### Cell Proliferation Assay

Cells were plated and grown to 70-90% confluence before transfection was performed in a 96-well plate. Post-transfection, the medium was refreshed every 48 hours and cells were cultured for an additional 72 hours. For the proliferation assay, the culture medium was mixed with CCK-8 solution in a 10:1 ratio. The original medium in the 96-well plate was carefully aspirated and the cells were washed twice with PBS. Each well was then filled with 100 µL of the medium-CCK-8 mixture. The plate was incubated for 2 hours in a cell culture incubator. Absorbance was measured at 450 nm. Cell viability was calculated using the formula: [OD (OE-CD24) - OD (blank)] / [OD (OE-control) - OD (blank)] × 100%.

### Apoptosis Assay

Apoptosis was assessed 72 hours post-transfection. Cells were detached using trypsin without EDTA. The cells were then centrifuged at 300g for 5 minutes at 4°C to facilitate collection. Following centrifugation, the cell pellet was washed twice with PBS. After discarding the supernatant, cells were resuspended in 100μL of 1× Binding Buffer. To stain the cells, 5μL of Annexin V-FITC and 10μL of PI Staining Solution were added to the cell suspension, followed by gentle mixing. The cell suspension was then incubated for 10-15 minutes at room temperature in the dark. After incubation, 400μL of diluted 1× Binding Buffer was added to the samples, which were then mixed well prior to flow cytometric analysis.

### Cell Transwell Assay

Seventy-two hours after seeding, the Transwell inserts were removed from the culture wells. The medium inside the inserts was discarded, and cells remaining inside the inserts were gently wiped off using cotton swabs moistened with PBS. To stain the migrated cells, 1 mL of crystal violet staining solution was added to each well of a clean 24-well plate, allowing the cells to stain for 3 to 10 minutes. After staining, the inserts were washed 2 to 3 times with PBS to remove excess stain. The washed inserts were then photographed to document the results of the cell migration assay.

### Wound-healing assay

Cells were transfected with plasmids for 24 hours, then scratched with a pipette tip to create a wound. The monolayer was washed once with PBS and then covered with complete culture medium. Initial wound images were captured immediately after washing, and the cells were incubated for an additional 72 hours to allow for migration and proliferation into the wound area.

### Statistical analysis

R software (version 4.3.1) was used for all statistical analyses. A statistically significant criterion of p < 0.05 was employed unless explicitly mentioned otherwise.

## Results

### 1.1 Screening of DEGs between BC and para-carcinoma tissues

The 765 diverse PCD-related genes were extracted from previous studies. Next, genes whose expression was 0 in half of the samples were removed during data processing. By comparing the expression levels of these genes between 1058 BC samples and 99 para-carcinoma samples in TCGA, 295 DEGs were identified. The volcano plot is displayed in Figure 1A. The PPI network is depicted in Figure 1B.

**Figure 1.**
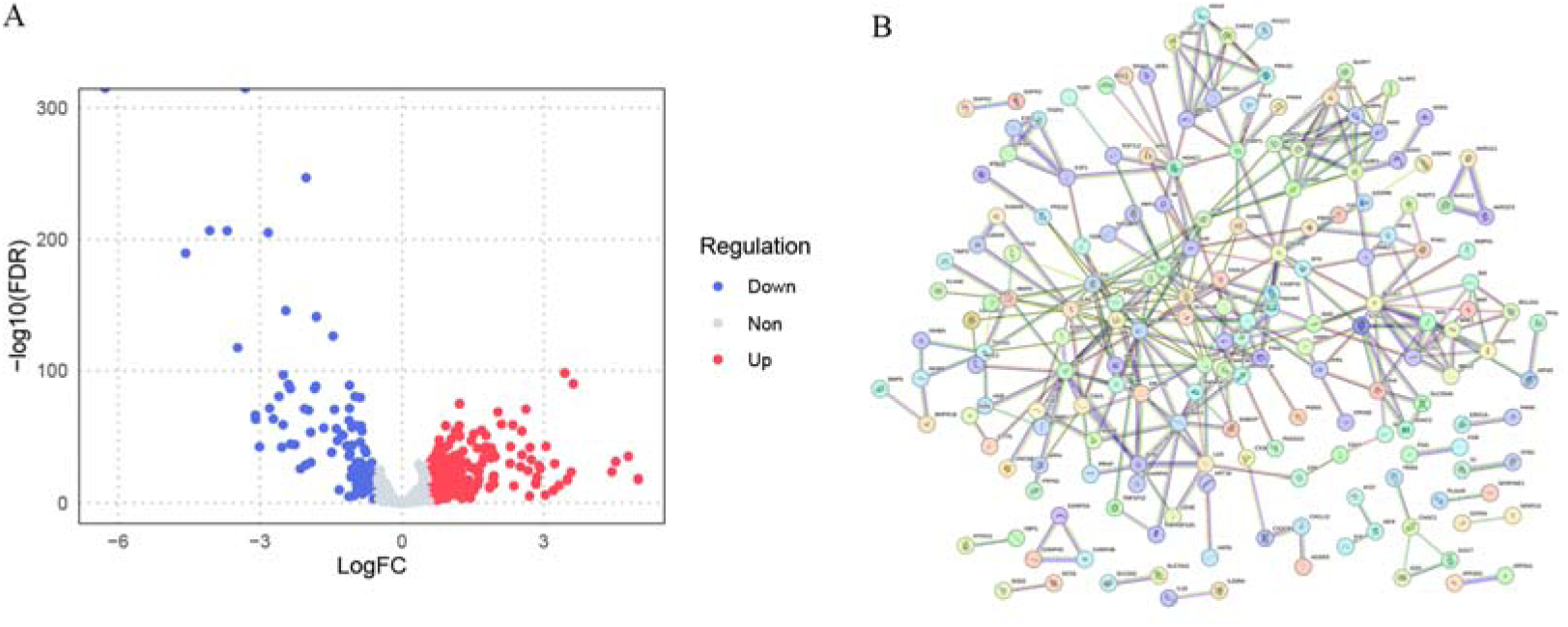
DEGs between BC and para-carcinoma tissues. (A) Volcano plot of 295 DEGs. (B) PPI network.

### 1.2 Univariate Cox analysis

Univariate Cox analysis was performed to screen survival-related genes among the 295 DEGs. This analysis resulted in the selection of 62 genes at a p-value of less than 0.03. To investigate the interactions among these 62 genes and assess their prognostic significance, we constructed a network (Figure 2).

**Figure 2.**
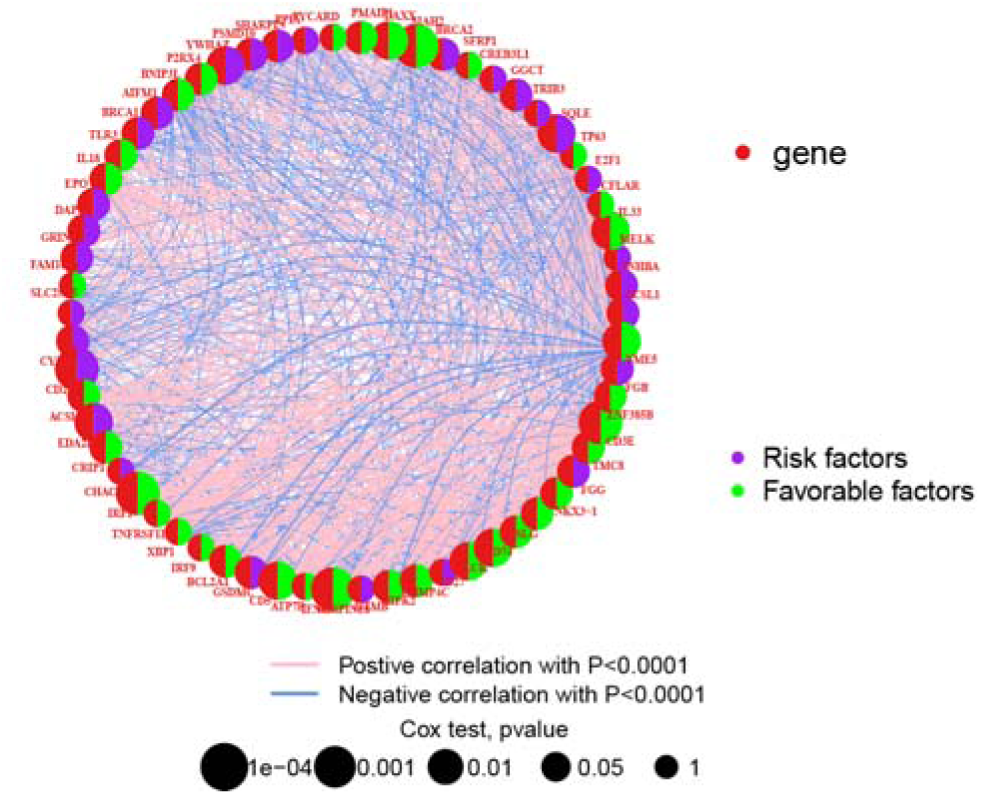
The interaction among 62 prognosis-related DEGs. Lines connecting genes represent their interactions, and the circle size reflects each regulator’s prognostic impact, scaled by its P-value. Green dots in the circle center indicate favorable factors for patient survival, while purple dots indicate risk factors.

### 1.3 Construction and validation of the prognostic model

We conducted LASSO regression analysis on the 62 OS-related DEGs, and the variability in the coefficients of these variables is depicted in Figure 3A. The optimal lambda value was determined based on the minimum partial likelihood deviance (Figure 3B). Subsequently, we constructed a Cox regression model. Ultimately, we identified 12 genes (CREB3L1, SFRP1, SHARPIN, AIFM1, IL-18, CD24, EDA2R, CRIP1, XBP1, BCL2A1, NKX3-1, and NME5). The Cox coefficients for the selected genes were derived through multivariate Cox proportional hazards regression analysis. RS can be calculated as follows: RS = (0.28815* CREB3L1) +(-0.11548* SFRP1) + (0.29269* SHARPIN) +(0.45416* AIFM1)+(-0.26994*IL-18)+(0.15480*CD24)+(0.36831* EDA2R)+(-0.32758* CRIP1) + (-0.33928*XBP1)+(-0.36601* BCL2A1)+(-0.12182* NKX3-1) +(-0.21151* NME5). RS for the testing cohort were computed using the established risk model.

**Figure 3.**
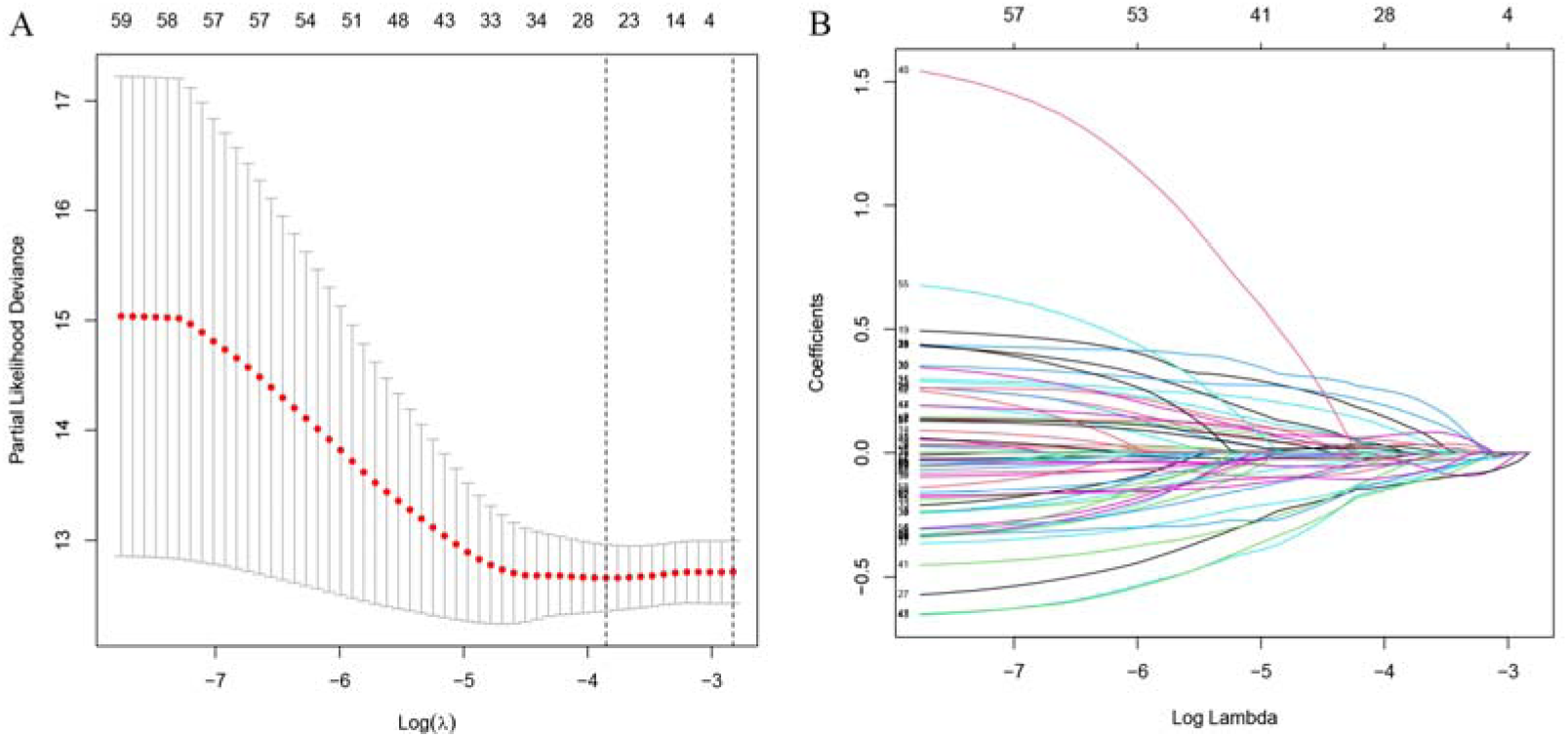
Construction of prognostic model in the training cohort. (A) LASSO regression of the 29 survival-related genes. (B) Cross-validation to fine-tune the selected parameter.

Consequently, patients in the training cohort were stratified into different RS subgroups based on the median RS (Figure 4A). Similarly, patients in the validation cohort were categorized into different subgroups using the same RS cutoff value as determined from the training cohort (Figure 4E). Patients with lower RS exhibited more favorable prognostic outcomes compared to those with higher RS (Figures 4B and 4F). The survival curves further illustrate that patients with low RS had superior survival rates in both the training and testing datasets(Figures 4C and 4G).

**Figure 4.**
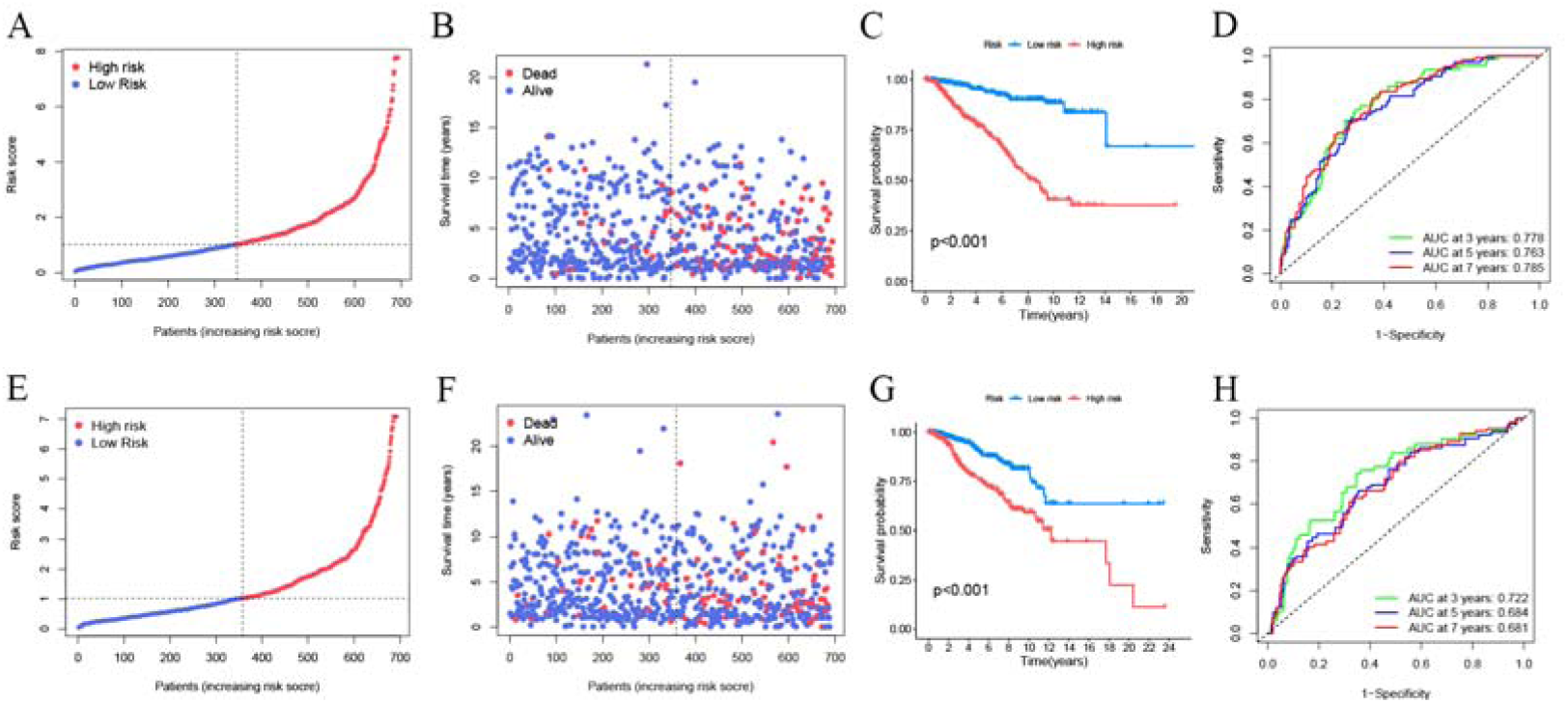
Validation of the model. (A) The distribution and cutoff value of RS in the training cohort. (B) The distribution and survival status of patients with different risk subgroups in the training cohort. (C) K-M curves for the training cohort. (D) ROC curves of training cohort. (E) The distribution of RS and their threshold value. (F) The distribution and survival status of patients with different risk subgroups in the testing cohort. (G) K-M curves for the testing cohort. (H) ROC curves of the testing cohort.

The AUC was calculated separately for 3-year survival (AUC = 0.778), 5-year survival (AUC = 0.763), and 7-year survival (AUC = 0.785) in the training cohort (Figure 4D). The AUC for RS at 1-year, 3-year, and 5-year OS in the testing group was 0.722, 0.684, and 0.681, correspondingly (Figure 4H). These results indicate that the prognostic prediction of the model demonstrated good efficiency.

### 1.4 Independent prognostic analysis

Multifactor Cox regression analysis was conducted, considering other clinicopathological factors such as gender, age, M, N, and T. Age, M, N, and the RS demonstrated independent prognostic significance (Figure 5A). This outcome affirms that, even when accounting for other potential confounding factors, the RS remains a valuable predictive indicator capable of predicting survival probabilities.

**Figure 5.**
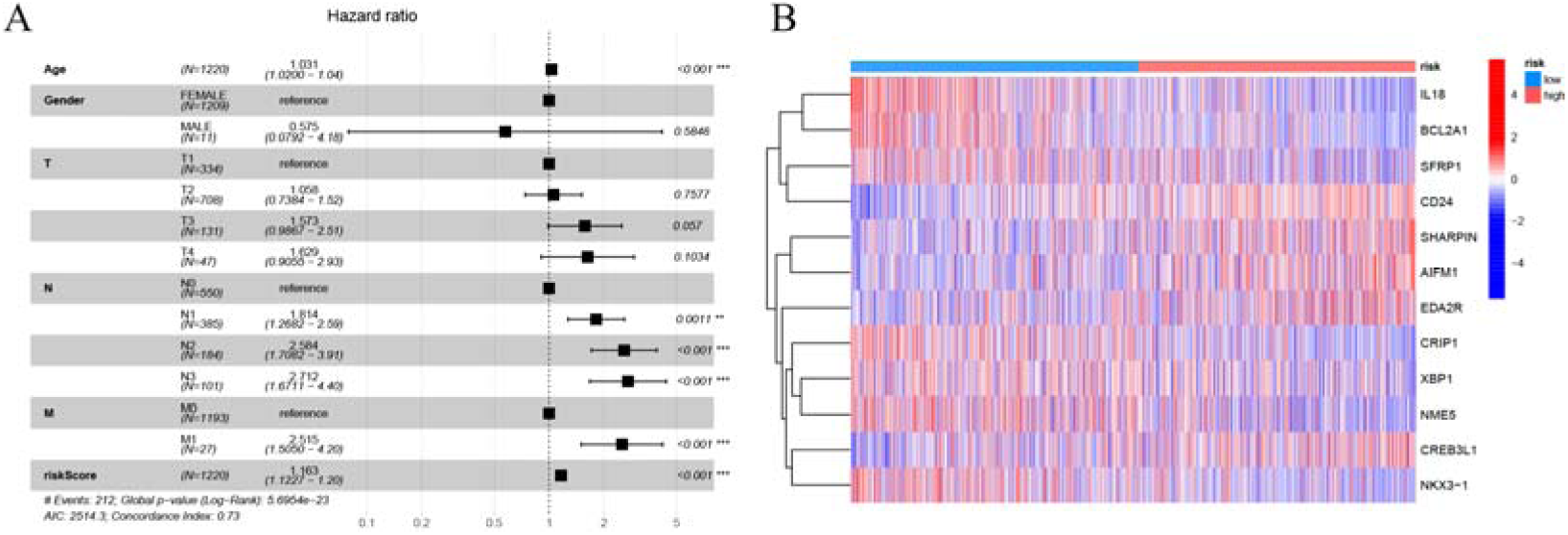
Independent prognostic analysis. (A) Multifactor analysis. *p < 0.05, **p < 0.01, and ***p < 0.001 were the displayed P values. (B) A heatmap of the gene expression from the model for different risk group patients.

### 1.5 Nomogram construction and validation

We have developed an innovative prognostic nomogram to offer a dependable and quantifiable method for predicting the survival of BC patients based on their RS and other clinicopathological factors. The nomogram assigns a corresponding score to each factor, and the cumulative score serves as a predictive tool (Figure 6A). To assess its accuracy, a calibration curve was generated (Figure 6B). The results indicate that the nomogram effectively predicts the overall survival probabilities. Additionally, cumulative hazard curves demonstrate that patient risk increases over time, with those possessing higher nomogram scores facing greater risks than those with lower scores (Figure 6C).

**Figure 6.**
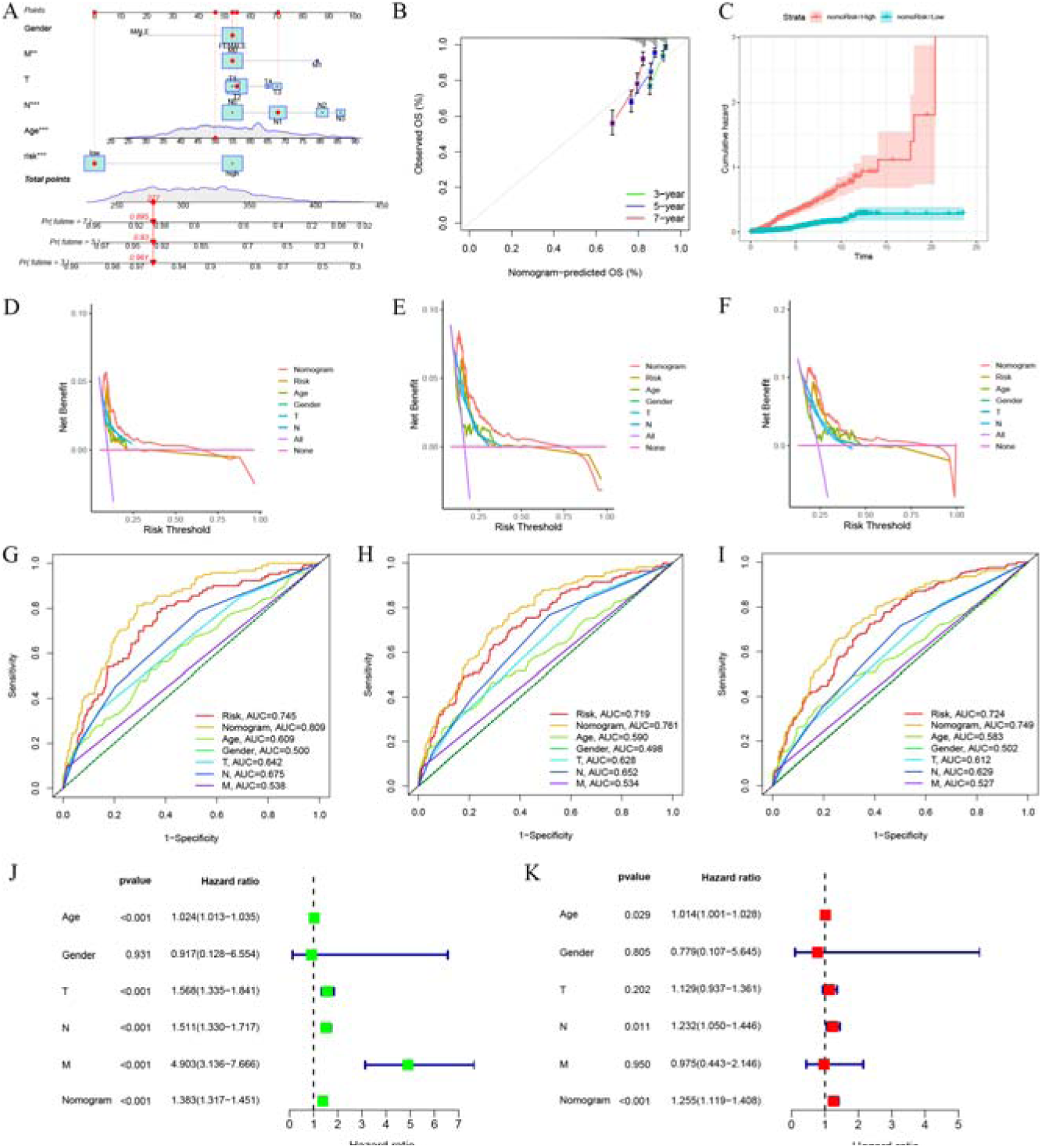
Nomogram construction and validation. (A) A nomogram was constructed to facilitate prediction. (B) Calibration curve of the nomogram. (C) Cumulative hazard curves. (D) 3-year DCA curves. (E) 5-year DCA curves. (F) 7-year DCA curve. (G) ROC curves of the nomogram and other factors for 3-year OS prediction. (H) ROC curves for 5-year OS prediction. (I) ROC curves for 7-year OS prediction. (J) Univariate independent prognostic analysis. (K) Multivariate independent prognostic analysis.

We further employed 3, 5, and 7-year DCA curves to illustrate the clinical utility of the nomogram (Figures 6D-F). By comparing the AUC values of the nomogram with other factors, the nomogram displayed superior predictive performance (Figures 6G-I). The result of the multivariate independent prognostic analysis confirmed that the nomogram remained a significant prognostic factor capable of predicting survival probability (Figures 6J and 6K).

### 1.6 Enrichment Analysis of the DEGs between different groups

To clarify the biological functions that underlie the prediction model, we extracted DEGs between different risk subgroups. Based on the identification of 144 DEGs, we conducted KEGG and GO enrichment analyses. The results of the GO enrichment analysis revealed that the DEGs were primarily associated with leukocyte-mediated immunity, lymphocyte-mediated immunity, regulation of T cell activation, and positive regulation of cell activation (Figure 7A). In terms of KEGG pathway analysis, the DEGs were predominantly linked to hematopoietic cell lineage, cell adhesion molecules, and Th1 and Th2 cell differentiation (Figure 7B).

**Figure 7.**
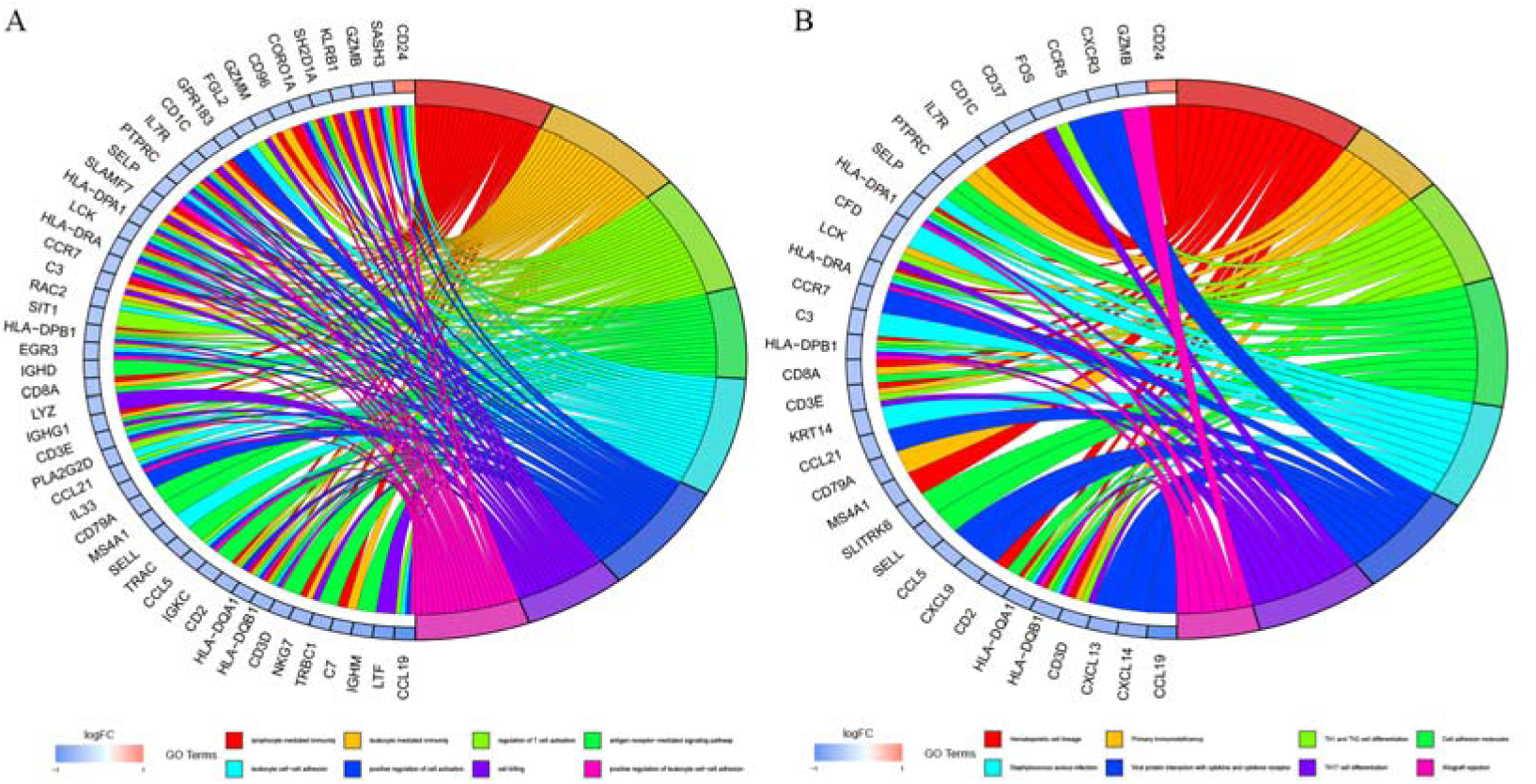
Enrichment analysis of the DEGs between different risk subgroups. (A) Chord plot for the enriched item by GO analysis. (B) Chord plot for the enriched item by KEGG analysis.

Furthermore, GSEA unveiled significant enrichment of pathways related to adaptive immune response, antigen receptor-mediated signaling pathway, B cell receptor signaling pathway, and lymphocyte-mediated immunity in the low-RS subgroup (Figure 8A). Conversely, the high-RS subgroup exhibited enrichment in specific pathways, including metaphase anaphase transition of the cell cycle, negative regulation of nuclear division, and regulation of chromosome segregation. (Figure 8B).

**Figure 8.**
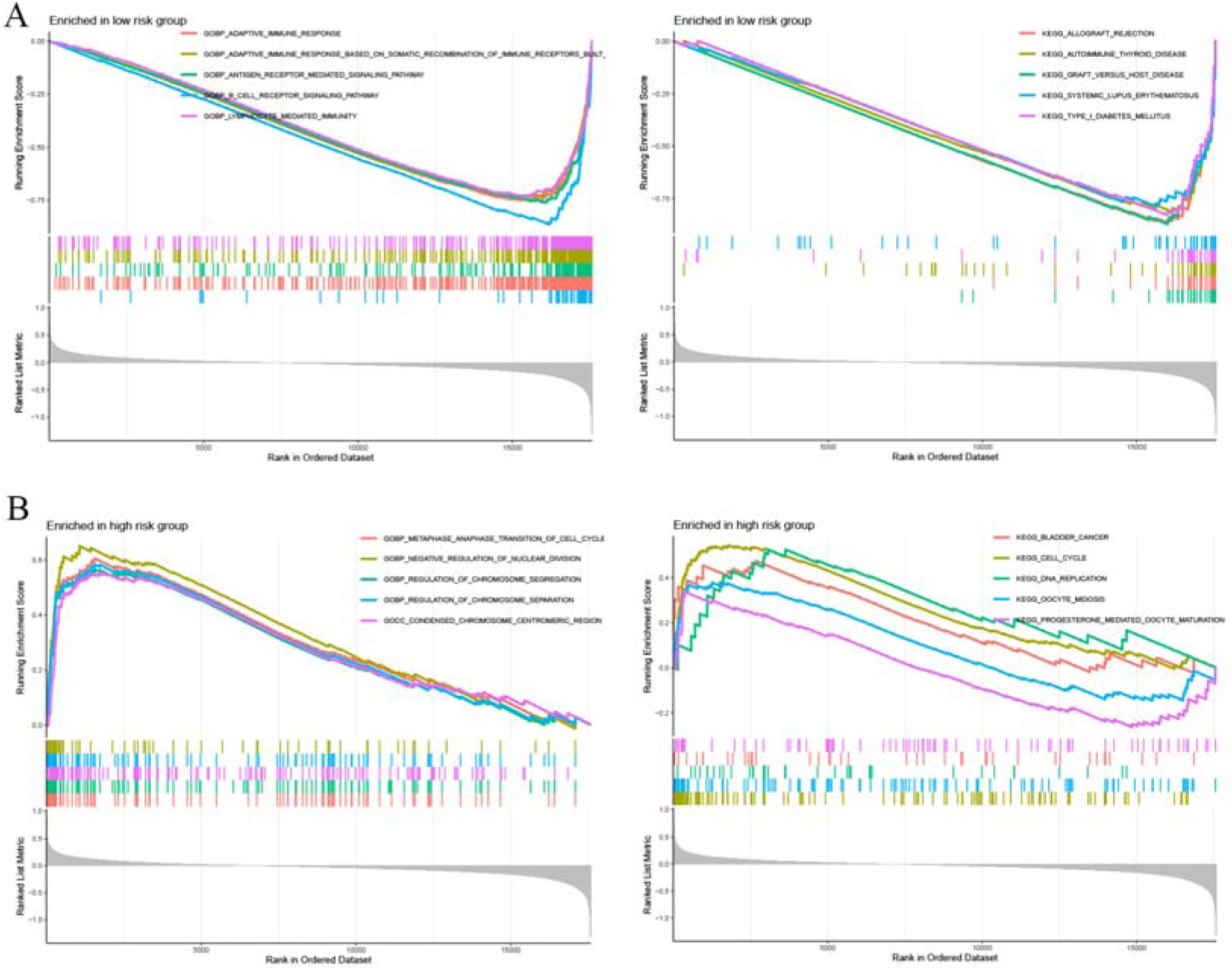
Gene Set Enrichment Analysis for the different risk subgroups. CC cellular component, BP biological process, MF molecular function.

### 1.7 Analysis of immune cell infiltration and TME

Using CIBERSORT, we assessed the immune infiltration levels of 22 cell types. We observed the immune infiltration levels of follicular helper T cells, CD8+ T cells, memory-activated CD4+ T cells, gamma delta T cells, and macrophage M1 in the low-RS subgroup were significantly higher than those in the high-risk subgroup. While, the high-RS subgroup exhibited higher levels of Tregs, macrophages M2, macrophages M0, and neutrophils (Figure 9A). The correlation among these immune cell populations is presented in Figure 9B. Additionally, we assessed the correlation between the immune cells and twelve genes (Figure 9C).

**Figure 9.**
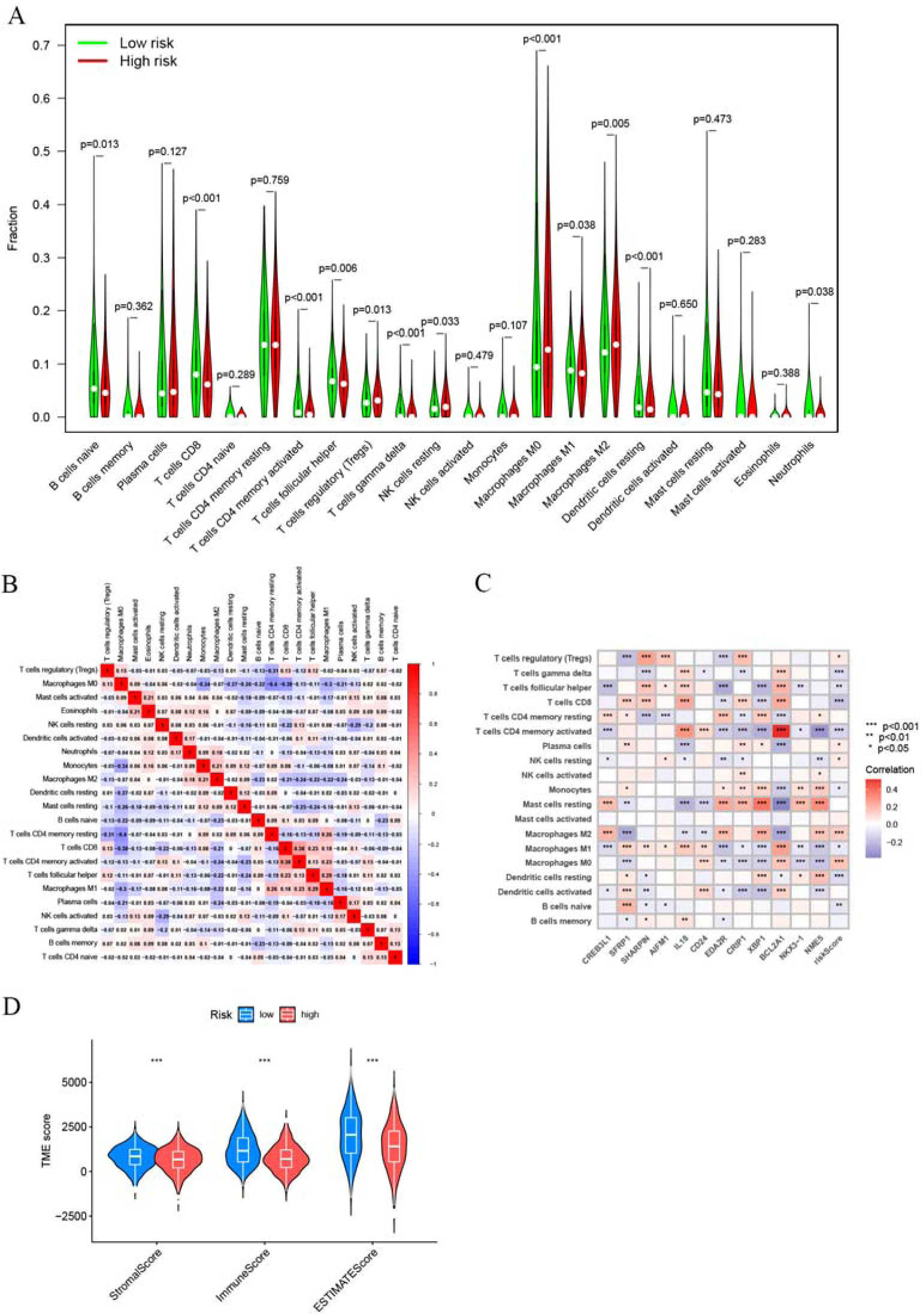
Analysis of immune cell infiltration and TME. (A) Differences in the expression of various immune cells in different risk subgroups. (B) The correlation of 22 types of immune cells (red signifies positive correlations, while blue represents negative correlations. The degree of association is shown by the color’s intensity). (C) Correlation between the abundance of immune cells and genes. (D) TME scores. The displayed p values were ***p < 0.001.

Furthermore, we assessed the TME scores which were notably higher in the low-RS group (Figure 9D). Higher ESTIMATE scores were associated with lower tumor purity. These findings imply that there was a considerable correlation between the immune state and RS.

### 1.8 Drug Sensitivity Analysis

Our data showed that the low-RS subgroup of BC patients had lower IC50 levels of axitinib, epirubicin, fulvestrant, and olaparib, suggesting that these patients exhibit greater sensitivity to these drugs (Figure 10A). Conversely, patients in the high-RS subgroup exhibited higher responsiveness to lapatinib, BI-2536, OSI-027, and SB505124 (Figure 10B). These results could be valuable in guiding medication selection for BC patients.

**Figure 10.**
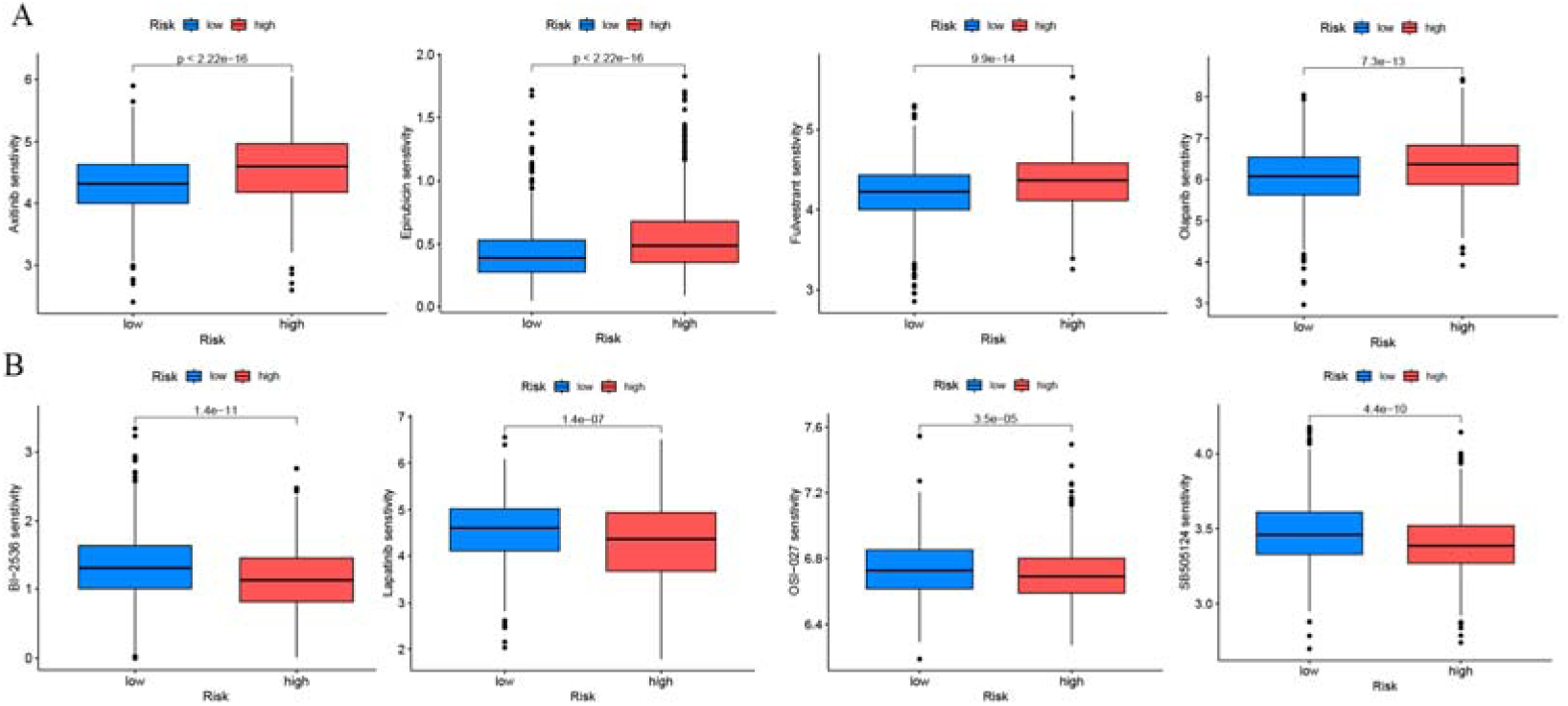
Drug sensitivity analysis. (A) Drugs with lower IC50 in the low-risk subgroup. (B) Drugs with lower IC50 in the high-risk subgroup.

### 1.9 Impact of CD24 Overexpression on Breast Cancer Cell Proliferation, Migration and Apoptosis

Our analysis resulted in a prognostic model centered on key genes implicated in cancer progression, among which CD24 was identified as a prime candidate for further investigation due to its unique expression pattern and underexplored role in BC. To explore the functional effects of CD24, we overexpressed it in BC cell line (Figure 11A-B) and performed various functional assays to examine its influence on cell behavior.

**Figure 11:**
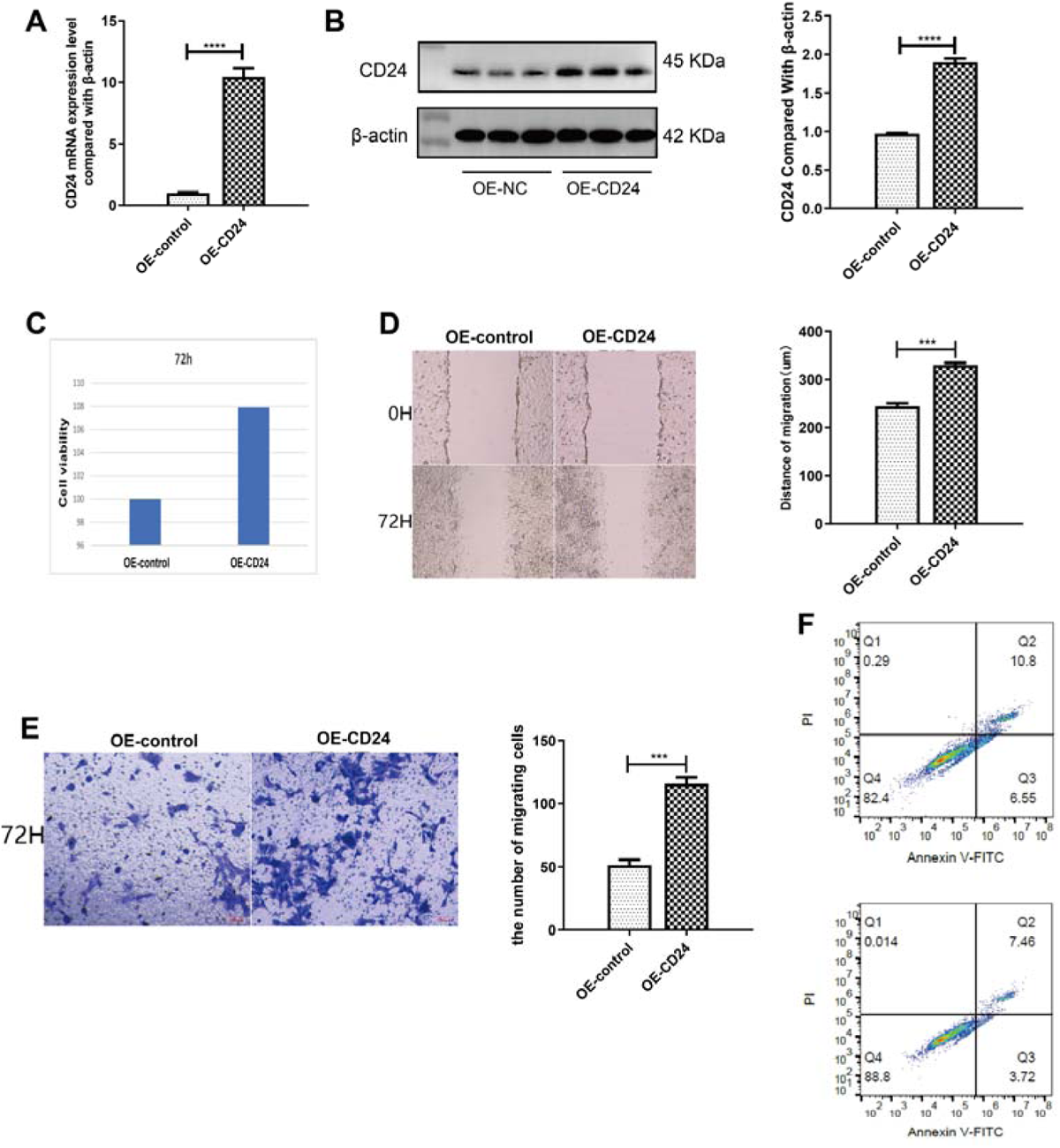
Impact of CD24 Overexpression on BC Cell Proliferation, Migration, and Apoptosis: (A) qPCR and (B) western blot analyses confirm CD24 overexpression in BC cells. (C) Proliferation curves showing the enhanced growth of BC cells with CD24 overexpression. (D) Scratch assay imagery and analysis illustrate the extended migration of BC cells overexpressing CD24 compared to control cells at 72 hours post-scratch, indicating increased migratory capability. (E) Transwell migration assays. (F) Apoptosis analysis via flow cytometry using Annexin V-FITC/PI staining reveals the distribution of cells across living (Q4), early apoptotic (Q3), late apoptotic (Q2), and necrotic (Q1) stages, the top panels represent the control group, while the bottom panels showcase the group with CD24 overexpression. Statistical significance denoted by asterisks in the bar graphs, with *** indicating P<0.001 and **** indicating P<0.0001.

The overexpression of CD24 resulted in a significant enhancement of cell proliferation, as evidenced by CCK8 assays, indicating increased proliferative capabilities in CD24-overexpressing cells (Figure 11C). Following this, in scratch assays, a notable acceleration in wound healing was observed in cells overexpressing CD24, demonstrating a substantial increase in migration speed compared to control groups in three separate trials (Figure 11D). This suggests CD24’s pivotal role in facilitating cell movement. Additionally, transwell migration assays provided further evidence of CD24’s function in promoting migration, with a significant uptick in the number of cells traversing the transwell membrane over 72 hours when compared to controls (Figure 11E). Moreover, apoptosis assays revealed a marked reduction in cell death among the CD24-overexpressing cells (Figure 11F), highlighting the gene’s contribution not only to cell migration but also to enhanced survival rates.

## Discussion

In recent years, multiple studies have highlighted the significance of PCD in the onset and advancement of various tumor types. PCD can impact the proliferation of tumor cells by influencing their immune microenvironment[26–30]. Therefore, it is crucial to investigate the interaction between PCD and the immune microenvironment to comprehend the mechanism underlying tumor development and to further enhance our understanding of BC therapy.

In our study, we developed a model incorporating twelve carefully selected genes (CREB3L1, SFRP1, SHARPIN, AIFM1, IL-18, CD24, EDA2R, CRIP1, XBP1, BCL2A1, NKX3-1, and NME5). Subsequently, we constructed a nomogram by integrating these genes with other clinicopathological factors. This nomogram was demonstrated to function as an independent prognostic tool, exhibiting a relatively robust predictive capability for the OS rates of BC patients. Notably, patients in the low-RS subgroup experienced longer survival times. GO enrichment analysis showed that the DEGs between the various subgroups were primarily linked to positive regulation of cell activation, lymphocyte-mediated immunity, leukocyte-mediated immunity, and regulation of T cell activation. Furthermore, GSEA indicated significant enrichment of pathways related to adaptive immune response, B cell receptor signaling pathway, and lymphocyte-mediated immunity in the low-RS subgroup. These results imply a connection between patient risk and immune response. Consequently, we conducted further comparisons of immune cell infiltration levels and the immune microenvironment between different subgroups. Research involving 12,439 BC patients suggests that heightened levels of CD8+ T cell infiltration in BC are linked to a significant reduction in the BC patient’s relative risk of mortality[31]. Tregs constitute a specialized subset of T cells that play a crucial role as mediators of immune tolerance. Through the inhibition of cytokine production and T-cell proliferation, they help to prevent autoimmunity. In the context of tumors, Tregs that infiltrate the tumor environment directly contribute to immune evasion and facilitate tumor growth[32, 33]. Follicular helper T cells aid in promoting adaptive immunity and are linked to a favorable prognosis in BC patients[34, 35]. In the context of tumor-associated macrophages, M1 macrophages exert an inhibitory effect on tumor growth, whereas M2 macrophages fulfill a pro-tumorigenic role by promoting angiogenesis, tumor development, and angiogenesis[36–38]. Our results align with the research findings. These results unveil the intricate interplay between the immune microenvironment and risk groups, providing valuable insights into potential therapeutic approaches for BC.

CAMP-responsive element-binding protein 3-like protein 1 (CREB3L1) operates downstream of PERK, particularly within the mesenchymal subtype of triple-negative tumors. Here, it exerts inhibitory effects on cancer cell invasion and metastasis, either through genetic or pharmacological means. Furthermore, CREB3L1 expression serves as a predictive marker for distant metastasis in patients with this specific tumor subtype[39]. Frequent alterations in CREB3L1 expression across various cancer types suggest its broader involvement in cancer progression and metastasis[40, 41]. Secreted Frizzled-related protein 1 (sFRP1) inhibits BC cell growth and metastasis by modulating the WNT pathway[42, 43]. SHANK-associated RH domain-interacting protein (SHARPIN) impacts tumorigenesis through ubiquitination processes and is linked to a poor prognosis of BC patients[44–46]. Diane Ojo et al. demonstrated that SHARPIN acts through the NF-κB and AKT pathways to promote tamoxifen resistance in BC cells[47]. Interleukin-18 (IL-18) has a controversial role in cancer[48–50]. In certain malignancies, CD24 is a highly expressed anti-phagocytic signal. ZBTB28 can prevent the growth of BC by downregulating the expression of CD24 and CD47, which in turn promotes macrophage phagocytosis[51, 52]. Cysteine-rich intestinal protein 1 (CRIP1) plays an important role in inhibiting BC proliferation and invasion[53]. X-box binding protein 1 (XBP1) and Hypoxia-inducing factor 1α (HIF1α) create a transcriptional complex that causes the carcinogenesis of triple-negative breast cancer (TNBC)[54]. BCL2A1, an anti-apoptotic BCL-2 family member, has been linked to poor prognosis in renal cancer patients undergoing immune checkpoint inhibitor therapy[55]. Studies have demonstrated that decreased BCL2A1 expression can enhance sensitivity to chemotherapeutic agents[56, 57]. In addition to acting as a tumor suppressor specific to the prostate, the androgen-regulated homeodomain transcription factor NKX3-1 is involved in the early stages of prostate development. Across various cancer types, NKX3-1 is associated with better patient prognosis[58, 59].

Our investigation has highlighted the significant impact of CD24 overexpression on cell proliferation, migration, and apoptosis in BC cell lines. Notably, the inclusion of CD24 in a prognostic model constructed from PCD-related genes underscores its pivotal role in cancer biology. This association suggests that CD24 may influence BC progression through mechanisms related to cell survival and death, aligning with its observed effects on apoptosis in our study.The differential expression of CD24, identified within our prognostic model of PCD-related genes, echoes findings from previous studies that have implicated CD24 in the regulation of cell death and survival pathways. For instance, research by Wang et al. demonstrated that CD24 can modulate apoptosis in gastric cancer through its interaction with signal transducer and activator of transcription 3 (STAT3)[60], highlighting its role in cell fate determination[61]. Our results extend this understanding to BC, suggesting that CD24 may similarly engage in key signaling pathways that promote cell survival and oppose programmed cell death, thereby facilitating tumor growth and metastasis. Furthermore, the correlation between CD24 overexpression and enhanced cell migration and proliferation observed in our study adds to the body of evidence supporting CD24 as a marker of aggressive cancer phenotypes[62].

However, there are several limitations in our research. Firstly, the conclusions drawn from statistical analyses are preliminary and await experimental validation. Secondly, to improve the robustness of our findings, a larger number of samples should be included.

In conclusion, our study has constructed a model and nomogram that offer improved prognostic predictions for BC patients. Furthermore, our research has provided insights into the BC tumor microenvironment and drug sensitivity, potentially opening new avenues for cancer treatment strategies.

## Ethical Approval

not applicable

## Availability of data and materials

The datasets generated and analyzed during the present study are available from the corresponding author on reasonable request.

Author Contribution Statement

Rihan Wu: Conducted research, gathered data, and wrote the introduction and literature review sections of the article.

Zirui Wang: Analyzed the data, interpreted the results, and wrote the methodology and findings sections of the article.

Chunhui Dong: Reviewed and edited the article for clarity, coherence, and accuracy, and provided valuable feedback for improvement.

Yihui Liu: Contributed insights, ideas, and expertise in the field, and helped shape the overall structure and argument of the article.

Ling Chen: Provided guidance, supervision, and coordination throughout the writing process, ensuring that the article met the highest standards of academic integrity and quality.

## Acknowledgement

NA

